# MIP3α-Rel_Mtb_ intranasal DNA vaccination induces reactive T-cell infiltration into the lungs in mice and macaques

**DOI:** 10.1101/2024.09.04.611263

**Authors:** James T. Gordy, Jean J. Zheng, Amanda R. Maxwell, Alannah D. Taylor, Styliani Karanika, Rowan E. Bates, Heemee Ton, Jacob Meza, Yangchen Li, Jiaqi Zhang, Petros C. Karakousis, Richard B. Markham

## Abstract

*Mycobacterium tuberculosis* (Mtb), the causative agent of tuberculosis (TB), is the leading cause of mortality due to a single infectious organism. While generally curable, TB requires a lengthy and complex antibiotic regimen, due in large part to bacteria that can shift to a persistent state in the presence of antibiotic pressure. Rel_Mtb_ is the primary enzyme regulating the stringent response, which contributes to the metabolic shift of Mtb to a persistent state. Targeting Rel_Mtb_ with a vaccine to eliminate persistent bacteria through the induction of Rel_Mtb_-specific T-cell immunity in combination with antibiotics to kill dividing bacteria has shown promise in model systems. In a mouse model of Mtb infection, a vaccine created by genetically fusing *rel*_*Mtb*_ to the chemokine macrophage inflammatory protein 3α (*MIP3*α), a ligand for the CC chemokine receptor type 6 (CCR6) present on immature dendritic cells, has been shown to enhance T-cell responses and accelerate eradication of infection in mouse models compared to a vaccine lacking the chemokine component. In this study, immunogenicity studies in the mouse and rhesus macaque models provide evidence that intranasal administrations of the DNA form of the MipRel vaccine led to enhanced lung infiltration of T cells after a series of immunizations. Furthermore, despite similar T-cell immunity seen in PBMCs between MipRel and Rel vaccinations, lung and bronchoalveolar lavage cell samples are more enriched for cytokine-secreting T cells in MipRel groups compared to Rel groups. We conclude that intranasal immunization with a MIP-3α fusion vaccine represents a novel strategy for use of a simple DNA vaccine formulation to elicit T-cell immune responses within the respiratory tract. That this formulation is immunogenic in a non-human primate model historically viewed as poorly responsive to DNA vaccines indicates the potential for clinical application in the treatment of Mtb infection, with possible application to other respiratory pathogens. Future studies will further characterize the protective effect of this vaccination platform.

## INTRODUCTION

*Mycobacterium tuberculosis* (Mtb), the causative agent of tuberculosis (TB), accounted for 10.6 million illnesses and 1.3 million deaths in 2022, making it the second leading infectious killer behind SARS-CoV-2 in 2022 [1]. While most TB infections are curable, current therapeutic guidelines require a six (or more)-month regimen consisting of isoniazid (INH) and at least three other drugs [1]. TB disease requires a lengthy therapy schedule because a proportion of the bacteria under conditions of stress can shift to a non-replicating, persistent state that is tolerant to INH and other antibiotics that target actively growing bacteria [2–7]. A key bacterial pathway for making and maintaining this switch to a persistent state is the stringent response, which is regulated by Rel_Mtb_, a (p)ppGpp synthase/hydrolase [8, 9]. Studies in settings of Rel_Mtb_ deficiency show that the lack of this protein results in reduced Mtb survival upon nutrient starvation, in animal models, and when exposed to INH [10–14]. Thus, Rel_Mtb_ is an attractive target for intervention, including in settings of multidrug resistance [9].

Previous work in our laboratories has yielded a therapeutic DNA vaccine construct targeting Rel_Mtb_ that provides additional efficacy when combined with a treatment that targets dividing Mtb, such as INH [15]. A modified vaccine fusing the Rel_Mtb_ to macrophage-inflammatory protein 3α (MIP3α, CCL20), which interacts with CCR6 on immature dendritic cells (iDCs) [16–23], enhances vaccine efficacy by specifically targeting vaccine antigen to iDCs, increasing DC activation, and enhancing subsequent T-cell activation. We have shown previously that the addition of MIP3α to the Rel_Mtb_ DNA vaccine (MipRel) enhances therapeutic responses in a murine model of chronic Mtb infection, and that the intranasal form of the vaccine showed the best anti-Mtb response and the lowest Mtb lung burden [24].

A similarly constructed vaccine with MIP3α fused to SARS-CoV-2 spike receptor binding domain (Mip-RBD) was administered intranasally and was found to elicit a robust anti-RBD T cell response [23]. Interestingly, when compared to control vaccines, one with MIP3α administered in the muscle and without the MIP3α given intranasally, it was discovered that all three were able to elicit similar levels of a systemic T-cell response by measure of antigen-specific splenic T cells able to produce Th1 cytokines. However, the Mip-RBD intranasal administration resulted in more overall T cells infiltrating into the lung environment, including significantly higher levels of T cells able to produce activation cytokines IFNγ and TNFα. The central hypothesis of this study is that the phenotype seen previously with the Mip-RBD vaccine is one that is likely universal with the MIP3α-antigen fusion DNA vaccine constructs, and that the MipRel DNA vaccine administered intranasally will likewise induce equivalent systemic responses with enhanced overall and reactive T-cell infiltration into the lungs, the critical site of Mtb infection.

## METHODS

### Vaccine Production

Rel and MipRel vaccines in mammalian expression system pSecTag2b are prepared as naked plasmid in 1xPBS, with same methodology as previously published [24]. In brief, DNA is housed in DH5-α (Thermo Fisher Scientific, Waltham, MA) *E. coli*, grown in large cultures, and extracted utilizing Qiagen’s (Germantown, MD) endotoxin-free kits. Prior to use, DNA is verified for purity, correctness, and concentration by insert sequencing (Johns Hopkins Synthesis and Sequencing Facility), nanodrop (Thermo Fisher Scientific, Waltham, MA) spectrophotometry, and agarose gel electrophoresis of undigested, single, and double digested samples. Prior to macaque immunizations and to avoid potential bacterial contamination, the vaccine DNA was passed through a 0.22 μm filter and the concentration was again verified by nanodrop.

### Mice

Mouse immunogenicity studies were performed primarily for a previous publication [24]. Data presented here utilized cryopreserved lung single cell suspensions from that study. In brief, vaporized isoflurane-anesthetized female C57BL/6 mice were intranasally inoculated with 200μg of vaccine plasmid DNA in 100μl of PBS (50μl administered per nostril). Mice were given three vaccinations at weekly intervals, and then 6 weeks post priming were euthanized, tissues were harvested, processed into single cell suspensions, and cryopreserved (Figure 1A).

**Figure 1:**
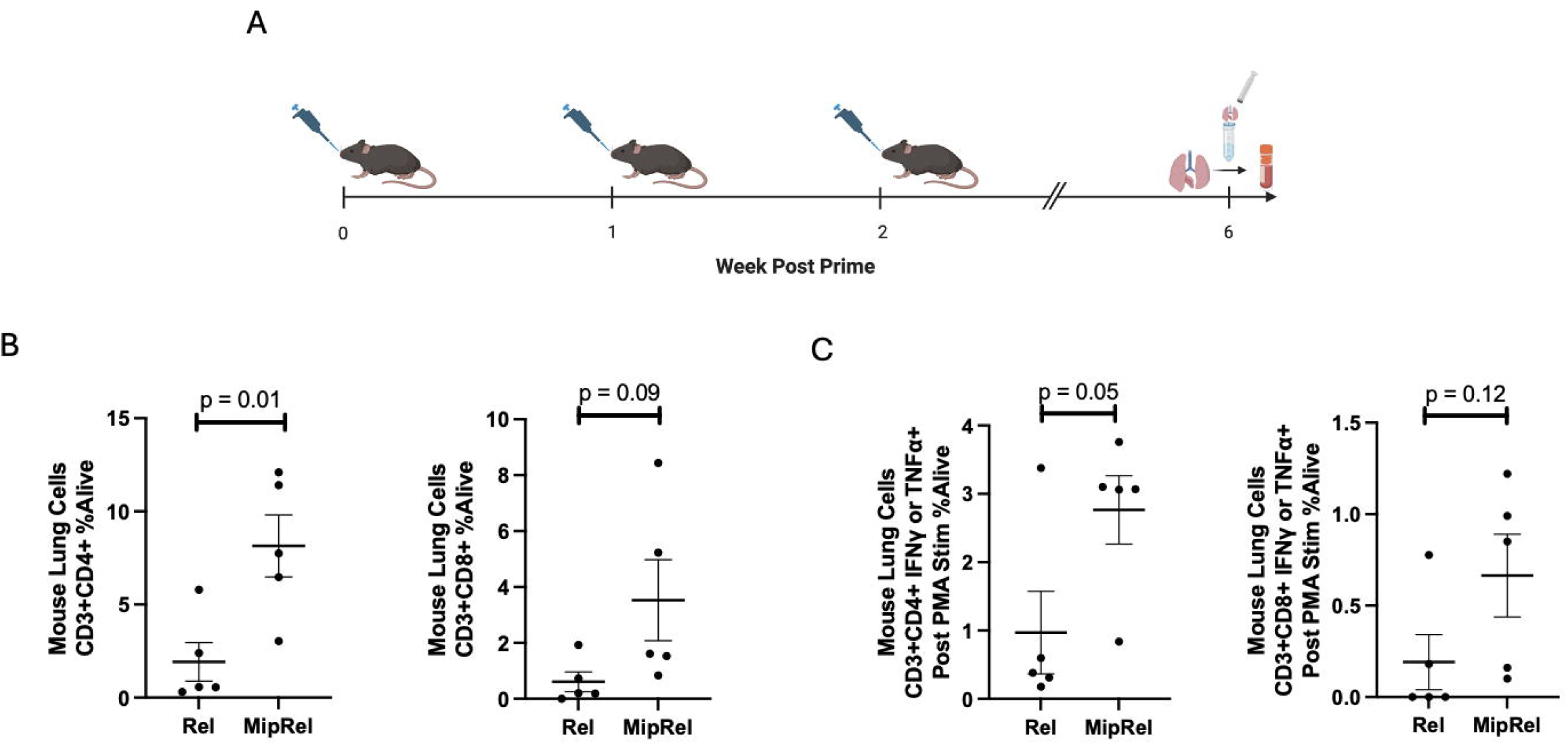
Mouse immunogenicity lung T cell infiltration data. A) Study outline of murine intranasal immunizations with tissue processing and cryopreservation at 6 weeks post prime. B) T cell proportions of all alive single cells in the homogenized lung samples. C) Proportion of lung T-cells secreting an activation cytokine after stimulation with PMA and ionomycin as a % of all alive single cells. Data are representative of one animal experiment with 5 mice per group, analyzed by Student’s T-Test with p-values labeled. Panel A created using Biorender.com.

### Macaques

1–3-year-old male and female rhesus macaques were kept in accordance to an approved JHU ACUC protocol. All animals were housed at the Johns Hopkins University Breeding Farm in singleL□species harem breeding groups or same sex juvenile/young adult groups. Animal enclosures consisted of runs with concrete flooring or raised corncrib cages. All animals had indoor and outdoor access. Animals were fed a standard commercial diet (rhesus macaques: 5049 FiberL□Plus Monkey Diet, LabDiet) and rotating food enrichment items including fresh fruits, vegetables, and dried fruit treats. Animals were provided water ad libitum. Annual colony health screening included intradermal tuberculin testing and serology for *Macacine herpesvirus* 1 (B virus), simian immunodeficiency virus, simian T□cell leukemia virus, and simian retrovirus. All animals were consistently negative on tuberculosis testing and viral serology. Macaques were randomized into vaccination groups with Rel group having 2M and 1F and MipRel 2F and 1M. Macaques received three immunizations administered at 3-week intervals. Prior to each immunization and one and three weeks after the final immunization, macaques underwent a bronchoalveoloar lavage (BAL) procedure and had blood drawn. Non-bronchoscopic BAL procedure [25]: Animals were fasted overnight and sedated with an intramuscular injection of 10 mg/kg ketamine HCl (Ketathesia, Henry Schein Animal Health) and 0.015 mg/kg dexmeditomidine hydrochloride (Dexmedesed, Dechra, Overland Park, KS). Once sedated, cetocaine spray was applied to the laryngeal folds of some animals to prevent laryngospasm and facilitate intubation. Animals were then be intubated with a sterile endotracheal tube to the approximate level of the thoracic inlet. Once intubated, animals were placed in left lateral recumbency, with the head and neck extended. 100% O2 (1 to 3 L/min) was administered to all animals immediately before and during the procedure via a nasal cannula attached to the endotracheal tube connector. Heart rate and peripheral oxygen saturation were continuously monitored via pulse oximetry. Depending on endotracheal tube size, an 8 or 10 french sterile, single-use pediatric suction catheter (Airlife Tri-Flo Suction Catheter with Control Port, Carefusion, Yorba Linda, CA) was inserted through the endotracheal tube connector and blindly passed through the trachea and into a mainstem bronchi. The catheter was passed until resistance was felt, indicating the catheter had wedged into a distal bronchus. For animals weighing 3 kg and above, 3 to 4 lavages of 11 mL each of sterile 0.9% saline were performed (33 to 44 mL total). For animals weighing less than 3 kg, 3 lavages of 6 mL each were performed (18 mL total). To perform the lavage, a 20 mL syringe containing the lavage fluid was attached to the end of the catheter and infused over 1 to 2 s. Immediately after the infusion, the aliquot was manually aspirated into the syringe using gentle pulsating suction. If needed, the catheter was moved slightly (up to 5 mm) during aspiration to maximize fluid return. This process was repeated for 2 to 3 more aliquots. Lavage fluid was immediately placed into 50 mL conical tubes containing R10 media and kept on ice. While sedated, the animals had 4ml of blood withdrawn into EDTA-coated tubes and kept on ice. On days that the animals require vaccination, after sampling and while remaining sedated, the animals were immunized intranasally. Vaccine was delivered as 500 μg plasmid DNA in 250 μl dripped into each nostril via pipette. Sedated macaques were monitored throughout sedation (respiration rate, mucous membrane color) until fully recovered. After the procedure, anesthesia was reversed with atipamazole hydrochloride (Antisedan, Zoetis, Kalamazoo, MI), and all animals were extubated and allowed to recover.

### Macaque Tissue Processing

Tissues were processed and cryopreserved as soon as possible after collection. Tissues were processed under Biosafety-Level 2 conditions. Dilutions and washes utilized Hanks Balanced Salt Solution (Thermo Fisher Scientific, Waltham, MA). Blood in EDTA tubes was separated into PBMC and plasma layers using Ficoll-Paque PLUS (Cytiva, Marlborough, MA) according to manufacturer’s instructions. The PBMC fraction was removed and washed. Red blood cells were lysed with ACK Lysis Buffer (Quality Biological Inc, Gaithersburg, MD) according to manufacturer’s instructions and was repeated for all samples if necessary to remove >95% of red color in the pellet. After washing the PBMC pellet, the cells were resuspended in freezing media (90% FBS [Corning, Corning, NY]; 10% DMSO) at approximately 1 million cells per ml and were aliquoted into approximately 6 cryovials each. The plasma fraction was centrifuged at 1500xg for 15 minutes to remove cellular debris and the supernatant was aliquoted and stored at -80°C. BAL samples were centrifuged at 300xg for 10 minutes. Supernatants were aliquoted and stored at -80°C. Cells were washed and red blood cells were lysed for all if blood contamination was present. Cells were resuspended in freezing media and aliquoted into 1-3 tubes. If cell count was greater than 1 million, then the cells were aliquoted into 3 tubes, 0.5ml each. If between 500,000 and 1 million, 2 tubes, and if less than 500,000, one tube. Both PBMC and BAL cells were cryopreserved in Mr. Frosty [Thermo Fisher Scientific, Waltham, MA] isopropranol cryopreservation chambers at -80°C for at least overnight, and then moved into -150°C for long term storage.

### Flow Cytometry

The protocol is the same for mouse and macaque samples, excepting the antibodies used. Cryopreserved cells were thawed quickly in 37°C water bath and diluted with warm complete media (CM) [RPMI 1640 with glutamine, 10% FBS, 1x penicillin/streptomycin, 20mM HEPES, 1% sodium pyruvate, and 1% non-essential amino acids]. Cells were centrifuged for 7 minutes, 250xg at room temp. Cells were resuspended in 0.2-1ml of warm CM and left to rest at 37°C 5%CO_2_ with vented tubes overnight. Phorbol myristate acetate (PMA) and ionomycin (Cell stimulation cocktail, Thermo Fisher Scientific, Waltham, MA) and brefeldin A (BioLegend, San Diego, CA) were added at manufacturer suggested dilutions to each well and incubated at 37°C for 4 hours. Staining was performed as previously described [23, 24]. Briefly, cells are washed with FACS Buffer (1xPBS, 0.5% BSA[Cell Signaling Technology, Danvers, MA]), stained with Live Dead Fixable Near-IR dye [Invitrogen, Waltham, MA], blocked with an Fc receptor blocker, stained for extracellular CD3, CD4, and CD8 markers, fixed (Fixation Buffer, BioLegend, San Diego, CA), permeabilized (Intracellular staining perm wash buffer, BioLegend, San Diego, CA), and stained for intracellular IFNγ, TNFα, and (macaque only) IL-17A markers. Antibodies are listed in Table 1. Samples were analyzed on an Attune™ NxT (Thermo Fisher Scientific, Waltham, MA) cytometer. FlowJo (FlowJo, LLC Ashland, OR) software was used for analysis. Full-minus-one (FMO) and unstimulated controls were utilized for gating. Gating strategies are provided in supplementary figure 1.

**Table 1:** Flow Cytometry Antibody Panels.

Flow Cytometry
| Table 1: Flow Cytometry Antibody Panels |  |  |  |  |
| --- | --- | --- | --- | --- |
| Target | Fluorophore | Clone | Dilution Factor | Company |
| Fixable Live Dead | Near-IR | L34994 | 2000 | Invitrogen |
| Macaque Markers |  |  |  |  |
| CD8a | APC | LT8 | 100 | Invitrogen |
| CD3 | FITC | FN18 | 100 | Invitrogen |
| CD4 | AF700 | OKT4 | 100 | Invitrogen |
| IFNγ | PE-Cy7 | 4S.B3 | 100 | Invitrogen |
| TNFα | Percp-Cy5.5 | MAb11 | 100 | Invitrogen |
| IL-17A | PE | eBio64CAP17 | 100 | Invitrogen |
| FcR-Block | | polyclonal | 10μg | Invitrogen |
| Mouse Markers |  |  |  |  |
| CD8a | AF700 | 53-6.7 | 200 | BioLegend |
| CD3 | PE-Cy7 | 17A2 | 100 | BioLegend |
| CD4 | FITC | GK1.5 | 100 | BioLegend |
| IFNγ | APC | XMG1.2 | 100 | BioLegend |
| TNFα | Percp-Cy5.5 | MP6-XT22 | 100 | BioLegend |
| FcR-Block | | S17011E | 2μg | BioLegend |

### Statistics

Mouse data were analyzed by Student’s T-test. Macaque data were analyzed by 2-Way repeated measures ANOVA with multiple comparisons utilizing uncorrected Fisher’s LSD. All midlines represent group mean and error bars the estimation of the standard error of the mean (SEM). For all tests, p ≤ 0.05 was considered to be significant and p-values are labeled on the graphs. Prism Graphpad 10 (San Diego, CA, USA) was used for all statistical analyses and figure generation.

## RESULTS

### Mouse immunogenicity model

Figure 1A outlines the model system. The animal study was performed primarily as a part of our previously published study [24]. Mice were immunized with Rel or MipRel vaccines intranasally thrice at one-week intervals and tissues harvested and cryopreserved at week 6. After thawing, resting, and stimulating with PMA and ionomycin, T-cell proportions and reactivity were assessed by intracellular staining flow cytometry. The full gating strategy is outlined in Supplementary Figure 1. T cells were defined as single, alive, lymphocytes by SSC vs FSC, CD3+ and then either CD4+ or CD8+. While PMA and ionomycin can decrease the signal of CD4 [26], the signal was distinguishable above background FMO controls. To determine overall proportion of T cells in the sample, the cells are reported here as a percentage of the first two gates that eliminate cell clumps and dead cells, denoted as %Alive. Figure 1B shows that in the homogenized lung samples, MipRel-vaccinated animals had more than 4 times the level of CD4+ T cells than Rel-vaccinated animals (means of 1.9 vs. 8.4; p = 0.01) and almost 6-fold higher levels of CD8+ T cells, albeit with more variability leading to a trend (means of 0.6 to 3.53; p = 0.09).

Next the levels of lung-infiltrating T cells able to produce effector cytokines IFNγ and TNFα were analyzed. The samples were limited so the most sensitive assay was utilized: stimulating with PMA and ionomycin to generate a statistic of all T cells capable of being activated in the lung environment. Figure 1C shows that the proportions of CD4 (left) and CD8 (right) T cells producing an effector cytokine was roughly tripled on average in both cases with the MipRel fusion vaccine compared to the vaccine expressing only Rel. CD4+ T cells increased from an average of 1% to 2.8% (p = 0.05) and CD8+ T cells increased from 0.19% to 0.66% (p = 0.12).

### Macaque immunogenicity model: T cells

Rel and MipRel vaccines were then tested in a nonhuman primate immunogenicity model. Vaccines were administered intranasally by pipetting, using a more clinically relevant timeline of 3 vaccinations at 3-week intervals. Rhesus macaques were followed serially with BAL and blood samples taken prior to first and three weeks post the final vaccinations (Figure 2A). BAL fluid (BALF) samples were separated into cellular and acellular fractions, and blood into PBMCs and plasma.

**Figure 2:**
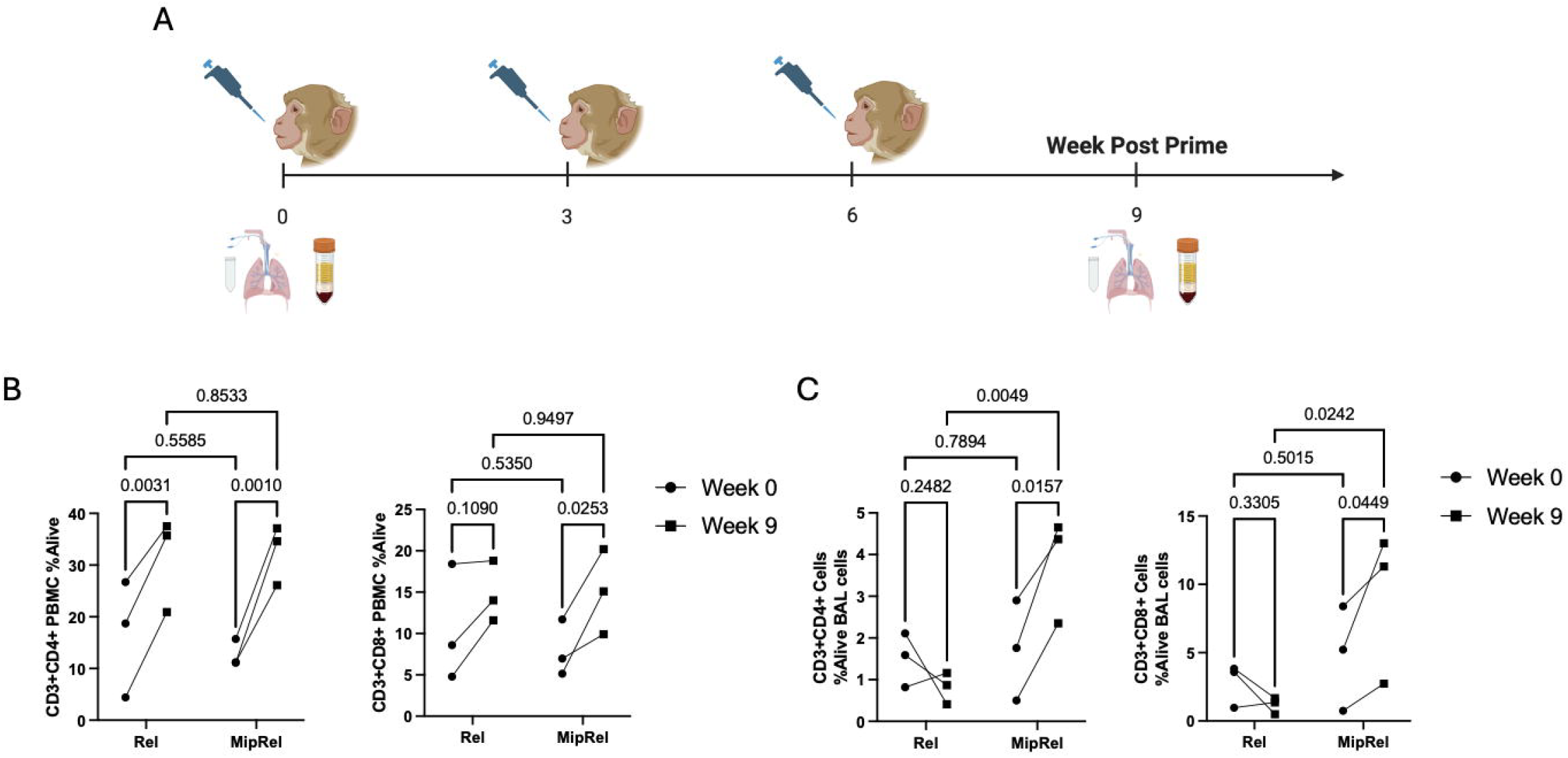
Macaque immunogenicity design and T cell levels. A) Rhesus macaques were intranasally inoculated with vaccine at three-week intervals. Prior to the first immunization and three weeks after the final immunization, broncho-alveolar lavage fluid and blood were collected. Lavage fluid was separated into cellular and acellular fractions and cryopreserved. Blood was separated into PBMCs and plasma and cryopreserved. All further data are from these samples. B) T cell proportions in PBMCs as a % of all alive single cells. C) T cell proportions in lavage cells as a % of all alive single cells. Data are representative of one animal experiment with 3 animals per group. Data were analyzed by repeated-measures two-way ANOVA, with p-values labeled. All further datasets are analyzed this way unless specified. Panel A created using Biorender.com.

Figure 2B analyzes the levels of T cells in macaque PBMCs for the two vaccines across the two timepoints. Overall CD4+ T-cell levels approximately doubled in their proportions in the PBMCs with both vaccinations comparing week 9 to 0, with all animals showing an individual increase (Rel Wk 0-9 p = 0.003; MipRel Wk 0-9 p = 0.001). Interestingly, comparisons across vaccine groups resulted in highly similar data at both time points. CD8+ T-cell levels had similar trends, although one Rel animal did not exhibit an appreciable increase over time, resulting in a nonsignificant trend between weeks 0-9 for Rel (p = .109). MipRel animals showed consistent increases of CD8+ T cells over time (p = 0.025). Again, levels across vaccination groups did not differ at either time point.

Figure 2C analyzes the levels of T cells in macaque BALF cells in a similar fashion. Intriguingly, the phenotype in the lung BALF cells differ from that in the PBMCs. Overall CD4+ and CD8+ T-cell proportions decreased slightly over time in the Rel-vaccinated animals. This is a significantly different situation as compared to MipRel vaccinated animals, which had a similar baseline to Rel animals but showed consistent increases in both CD4+ (p = 0.016 Wk 0-9) and CD8+ (p = 0.045 Wk 0-9) T cells that resulted in differences between vaccination groups (CD4+ p = 0.005; CD8+ p = 0.024).

### Stimulation of macaque PBMCs

The PBMCs were further analyzed for their ability to produce cytokines after stimulation with PMA and ionomycin. Antigen-specific stimulation analysis of PBMCs and BALF cells will be presented in a separate manuscript (manuscript in preparation). IFNγ and TNFα production by CD4+ T cells were analyzed in Figures 3A-B. Consistently, all animals across both vaccination groups showed an increase in cytokines from CD4+ T cells between weeks 0 and 9, but neither time point provided a difference between vaccinations. CD8+ T cells had more variability over time leading to non-significant trends with both cytokines (Figure 3 C-D). MipRel-vaccinated animals showed more consistent increases across individual animals (3/3) as compared to the Rel animals (2/3) for both CD8+ cytokines. IL-17A levels were measured but were low and therefore the data are not shown. Across the two groups, the datasets are highly similar at both timepoints.

**Figure 3:**
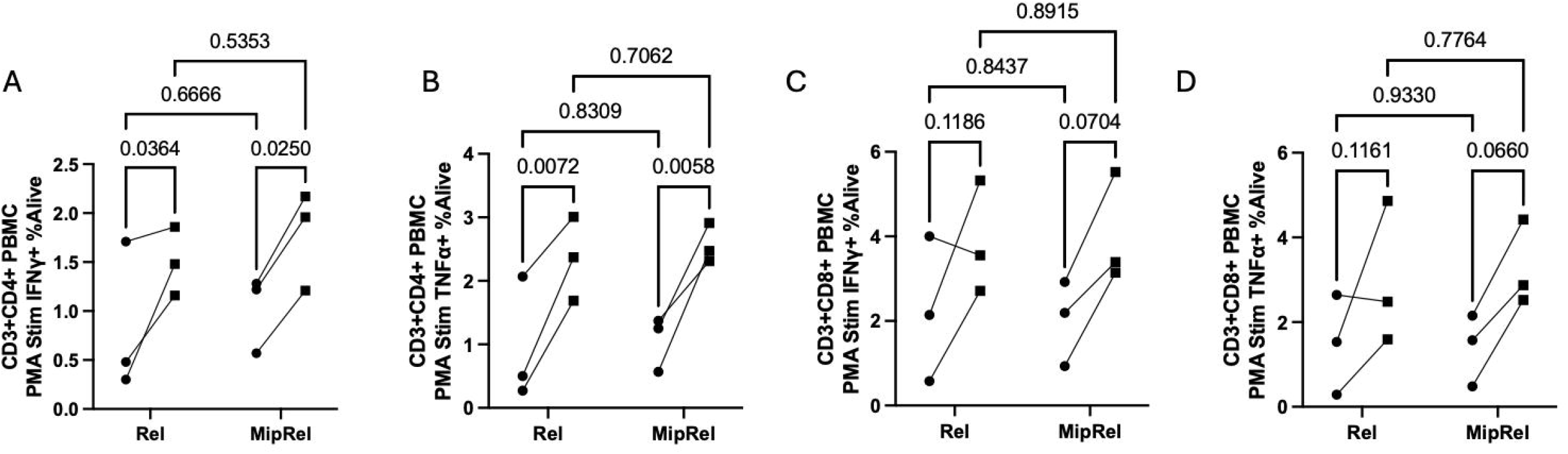
Stimulation of macaque PBMCs with PMA and ionomycin. Cryopreserved macaque PBMCs were quick-thawed, rested at 37°C 5%CO_2_ overnight, stimulated with PMA and ionomycin in the presence of Brefeldin A for 4 hours, stained extracellularly for CD3, CD4, CD8, and Live/Dead and Intracellularly with IFNγ, TNFα, and IL-17. A) CD4+ T cells producing IFNγ. B) CD4+ T cells producing TNFα. C) CD8+ T cells producing IFNγ. D) CD8+ T cells producing TNFα. In all cases, circles represent week 0 and squares week 9.

### Stimulation of macaque lavage cells

BALF cells were likewise analyzed for their ability to produce cytokines post-PMA and ionomycin stimulation. Across both CD4+ and CD8+ T cells and all three cytokines analyzed, Rel-immunized animals showed no increases above baseline, in contrast to the data seen in PBMCs. Levels of reactive T cells in almost all cases increased significantly from weeks 0 to 9 in MipRel-vaccinated animals. In CD4+ T cells, MipRel-vaccinated animals had increases from 0.25% to 1.83% of cells producing IFNγ (p = 0.007 Wk 0-9; p = 0.001 Rel-MipRel wk9) (Figure 4A) and from 0.23% to 1.69% producing TNFα (p = 0.012 Wk 0-9; p = 0.002 Rel-MipRel Wk9) (Figure 4B). In both cases, all three animals showed similar increases. IL-17A production was not uniform, with 2/3 MipRel-vaccinated animals increasing over time as opposed to 1/3 Rel animals (Figure 4C). The phenotypes of CD8+ T cells were similar to those of the CD4+ T cells. MipRel-vaccinated animals showed uniform increases over time in IFNγ-producing cells (1.08% to 4.36%) and TNFα-producing cells (0.60% to 3.1%), leading to significance across time (p = 0.02 and 0.03) and across vaccination groups (p = 0.01 and p = 0.01) for IFNγ and TNFα-producing cells, respectively (Figure 4D-E). While all three animals in the MipRel group showed increases in IL17A production over time in the CD8+ T cells (Figure 4F), the variability led to trends across time (p = 0.088) and across groups at week 9 (p = 0.07). Importantly, in Figures 4 G-H, T cells able to simultaneously secrete two cytokines were analyzed. The phenotypes seen in single-cytokine analysis hold true when looking at double cytokine production. Rel vaccinated animals remain at baseline over time while MipRel-vaccinated animals showed significant increases for both CD4+ and CD8+ T cells over time (CD4: 0.11% to 1.17%, p = 0.012; CD8: 0.37% to 1.83%, p = 0.049) and compared to Rel at week 9 (CD4+ p = 0.002; CD8+ p = 0.016). Across all panels both Rel-vaccinated and MipRel-vaccinated animals had similar levels at baseline.

**Figure 4:**
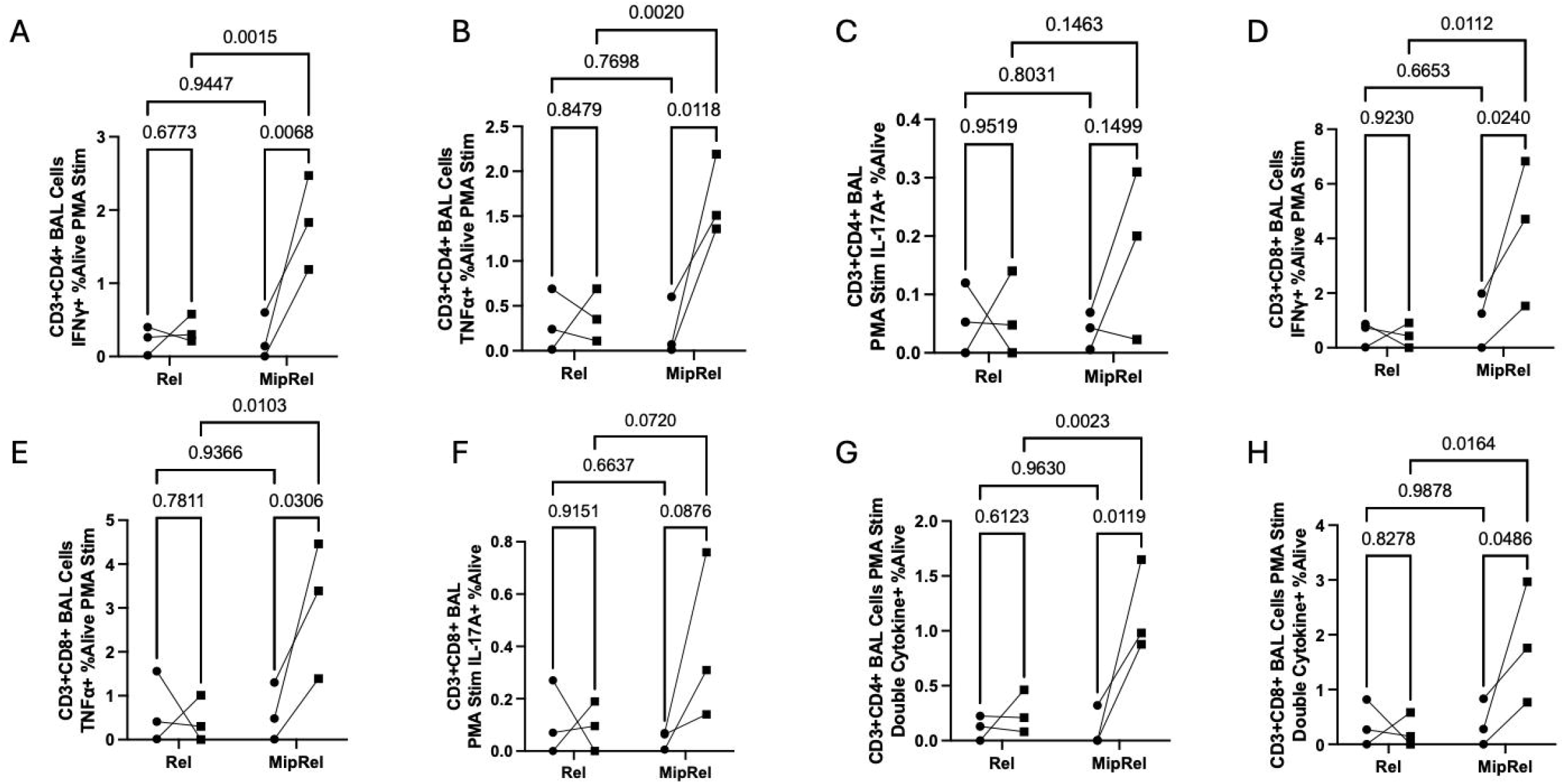
Stimulation of macaque lavage cells with PMA and ionomycin. Cells were treated the same way as PBMCs previously. A-C analyze CD4+ T cells and D-F CD8+ T cells, for each series the graphs analyze production of IFNγ, TNFα, and IL-17A respectively in that order. In all cases, circles represent week 0 and squares week 9. G) CD4+ and H) CD8+ T cells producing any combination of 2 aforementioned cytokines.

## DISCUSSION

In SARS-CoV-2 [23] and melanoma [18, 27] models, studies have shown that MIP3α fusion vaccinations can affect T-cell infiltration into tissue sites of interest. The SARS-CoV-2 vaccine in particular provided evidence that, in an intranasal setting, MIP3α presence in the vaccine can lead to enhanced levels of T cells infiltrating into the lung, even in the absence of measurable differences in splenocytes [23]. In this study, we show that three to four weeks after a three-vaccination series in two animal systems, mouse and rhesus macaque, intranasal administrations of a naked DNA vaccine encoding MIP3α fused to the Mtb stringent-response regulation gene Rel_Mtb_ (MipRel) increases the proportions of T cells present in the lung, both overall and those producing cytokines, compared to a Rel_Mtb_ vaccine without MIP3α. Both vaccines have shown in previous work in a mouse model to have efficacy against chronic TB, with the MIP3α fusion vaccine providing marked reduction in lung bacterial burden in chronically Mtb-infected mice and enhancements in antigen-specific T cells as measured by percent of T cells [15, 24]. This is the first report to show that cytokine-secreting T-cell levels in the lung are enhanced with the MipRel vaccine when analyzed by a percentage of all cells.

While Rel_Mtb_-antigen specific responses are important to this vaccine system, with limited macaque sampling, it was essential to use an assay with the highest sensitivity to glean T-cell responsiveness comparing the two vaccine types. Antigen-specific stimulation was conducted, and the MipRel elicitation of a specific T-cell response will be included in a manuscript in preparation. The assay was not sensitive enough, however, to detect significant differences between MipRel and Rel vaccinations with only 3 animals per group (data not shown), although MipRel trended higher with 3 out of 3 animals showing lung responses as compared to 1 out of 3 in the Rel-vaccinated group. Further, concerns have been raised about reduced assay sensitivity in BALF cells due to the complexity and length of time of the assays, as well as the potentially altered ability of alveolar macrophages to act as effective antigen presenting cells for Th1 cytokines. PMA and ionomycin is an appropriate alternative for overall T-cell responses in the lung environment [28]. Additionally, multi-cytokine responses after PMA and ionomycin stimulation have been shown to be clinically correlated with less severe TB disease [29].

Here, it has been shown that in the mouse model the intranasal MipRel vaccine leads to a greater influx of T cells of both CD4+ and CD8+ types into the lungs compared to the Rel vaccine, and that cytokine-producing cells after PMA and ionomycin stimulation are also increased in the lungs (Figure 1). Intranasal vaccination in rhesus macaques showed analogous responses, with increased levels of T cells in the lung, as well as increased levels in PBMCs (Figure 2). PMA and ionomycin stimulation of the PBMCs showed that both Rel and MipRel vaccines elicited responses over time. Interestingly, the PBMC responses induced by the Rel and MipRel vaccines were very similar, suggesting that both vaccines are able to elicit systemic T-cell responses to similar degrees (Figure 3). However, analysis of BALF cells painted a much different picture. MipRel-vaccinated animals had significantly higher levels of most (6/8) combinations of CD4+ or CD8+ T cells producing IFNγ, TNFα, IL17, or any two cytokines as compared to Rel-vaccinated animals at week 9 (Figure 4). Importantly, both vaccine groups had similar cytokine levels pre-vaccination, so differences cannot be attributed to baseline heterogeneity.

Based on the data presented here and previously [23], we can hypothesize that when administered intranasally, a primary mechanism of the Mip-antigen vaccine response is greater infiltration of T cells into the lung environment, either directly by interaction of MIP3α to CCR6, known to be present on memory T cells [30] or indirectly by other mechanisms. Future studies will analyze this phenomenon more thoroughly to better understand the dynamics of the increased cell population. The specific immunophenotyping of the recruited T cells utilizing more complex flow cytometry panels, with a focus on activation, effector/memory, and trafficking markers will be undertaken. Additionally, more thorough analysis of other cell types present, including dendritic cells, macrophages, B cells, and others will be performed to better understand the immune mechanisms associated with the enhanced responses that were observed.

## ACKNOWLEDGEMENTS

We would like to thank the JHU research animal resources veterinary post-doctoral fellows and the JHU breeding farm staff as well as the caretakers in the mouse facilities. We would like to thank Dr. Prakash Srinivasan, the Johns Hopkins Malaria Research Institute, and the Department of Molecular Microbiology and Immunology for Attune™ NxT flow cytometry support.

## AUTHOR CONTRIBUTIONS

JTG led the macaque study including vaccine production and quality control efforts, coordination with veterinary staff, tissue processing and cryopreservation, t-cell stimulation flow cytometry assays, data analysis, and manuscript writing. JJZ performed the stimulation and flow cytometry analysis of the mouse samples and assisted with flow cytometry analysis of the macaque samples and figure creation. ARM was responsible for macaque immunization, sample collection, and animal management. SK performed the mouse study. REB, ADT, YL, JM, HT, JJZ, and JZ assisted with macaque sample processing and cryopreservation. ADT, REB, JM, and YL assisted with vaccine production and quality control. RBM and PCK contributed to the conception and design of the experiments and the writing of the manuscript. All authors approved of the final draft.

## FUNDING SOURCES

This work was supported by NIH grants R01AI148710 to PK and RM and K24AI143447 to PK

## COMPETING INTERESTS

Authors JTG, SK, RBM, and PCK are inventors on pending patent: PCT/US2023/065584. All other authors declare no competing interests.

## DATA AVAILABILITY

Processed data are provided in the submission and are available upon reasonable request.

## FIGURE LEGENDS

<u>Supplementary Figure</u>: Gating Strategy using a representative macaque PBMC sample. The Mouse experiment was gated the same way except for IL17, which was not included in the panel.

