## Supplementary figures and images for "MIP3α-Rel_Mtb_ intranasal DNA vaccination induces reactive T-cell infiltration into the lungs in mice and macaques"

### Supplemental Figure 1

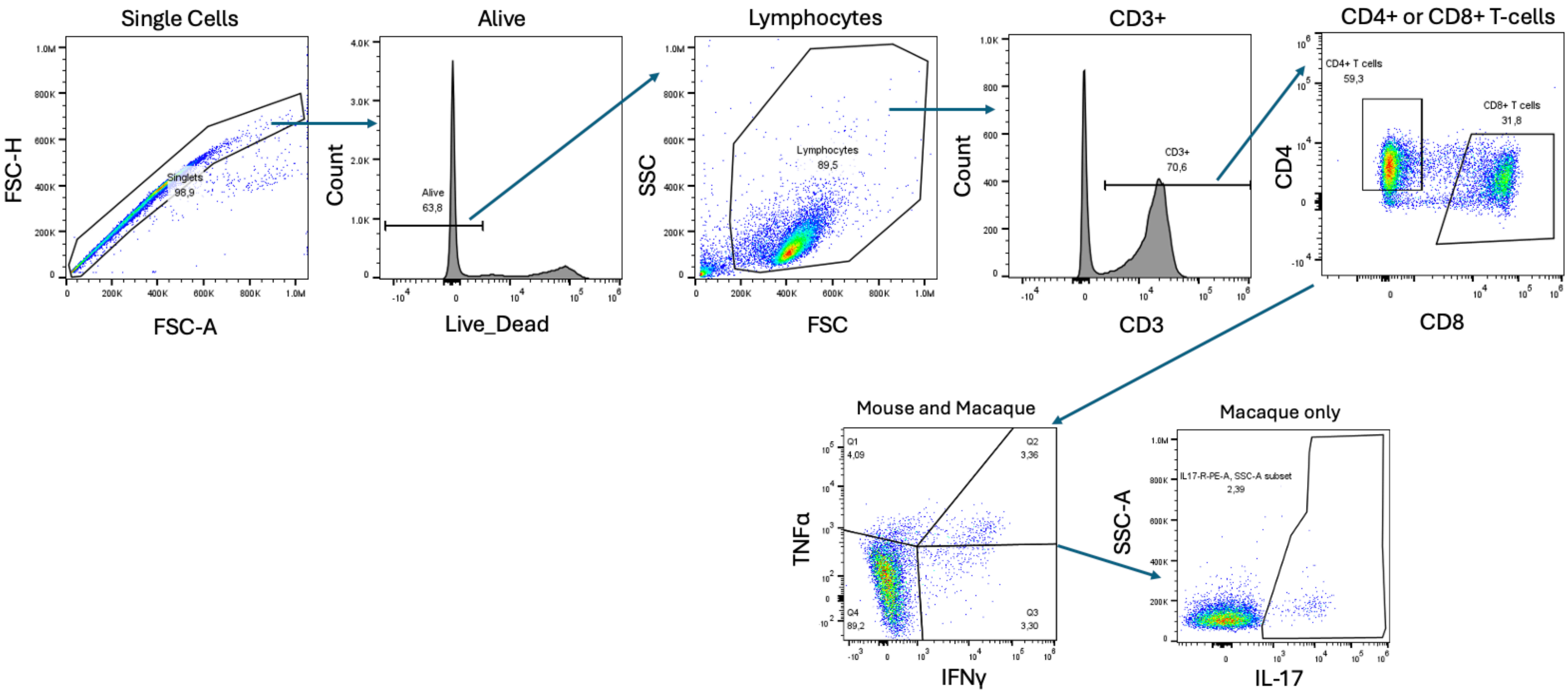
